# Protein kinase A catalytic-α and catalytic-β proteins have non-redundant functions

**DOI:** 10.1101/2020.07.01.182691

**Authors:** Viswanathan Raghuram, Karim Salhadar, Kavee Limbutara, Euijung Park, Chin-Rang Yang, Mark A. Knepper

## Abstract

Vasopressin regulates osmotic water transport in the renal collecting duct by PKA-mediated control of the water channel aquaporin-2 (AQP2). Collecting duct principal cells express two seemingly redundant PKA catalytic subunits, PKA catalytic α (PKA-Cα) and PKA catalytic β (PKA-Cβ). To identify the roles of these two protein kinases, we carried out deep phosphoproteomic analysis in cultured mpkCCD cells in which either PKA-Cα or PKA-Cβ was deleted using CRISPR-Cas9-based genome editing. Controls were cells carried through the genome editing procedure, but without deletion of PKA. TMT mass tagging was used for protein mass spectrometric quantification. Of the 4635 phosphopeptides that were quantified 67 were significantly altered in abundance with PKA-Cα deletion, while 21 were significantly altered in abundance with PKA-Cβ deletion. However, only four sites were changed in both. The target proteins identified in PKA-Cα-null cells were largely associated with cell membranes and membrane vesicles, while target proteins in the PKA-Cβ-null cells were largely associated with the actin cytoskeleton and cell junctions. In contrast, in vitro incubation of mpkCCD proteins with recombinant PKA-Cα and PKA-Cβ resulted in virtually identical phosphorylation changes. In addition, analysis of total protein abundances in the in vivo samples showed that PKA-Cα deletion resulted in a near disappearance of AQP2 protein, while PKA-Cβ deletion did not decrease AQP2 abundance. We conclude that PKA-Cα and PKA-Cβ serve substantially different functions in renal collecting duct cells and that differences in phosphorylation targets may be due to differences in protein interactions, e.g. mediated by AKAP, C-KAP or PDZ binding.

## INTRODUCTION

Vasopressin’s actions in collecting duct principal cells (23) are mediated by a G-protein coupled receptor (GPCR) V2R (Gene symbol: *Avpr2*), which couples to the heterotrimeric G-protein stimulatory α subunit (G_s_α) with activation of adenylyl cyclase 6 (22). Increased intracellular cyclic AMP levels result in physiological effects in collecting duct principal cells largely through activation of PKA (4, 5, 8, 11, 12, 17, 19–21) . Two of the most important physiological end effects are: 1) membrane trafficking changes that increase the abundance of the water channel protein aquaporin-2 (AQP2) in the plasma membrane (18) and 2) increased transcription of the *Aqp2* gene (7, 14, 24), both of which contribute to vasopressin induced increases in osmotic water transport across the collecting duct epithelium. Protein Kinase A (PKA) is a widely studied protein that has been viewed by most investigators as a single entity, although its catalytic subunits are coded in mammalian genomes by two separate genes, PKA catalytic α (PKA-Cα [Gene symbol: *Prkaca*]) and PKA catalytic β (PKA-Cβ [Gene symbol: *Prkacb*]). At an amino-acid level, the two are 91 percent identical and the catalytic domains are very similar. (Footnote: A third entity PKA catalytic γ, is not widely expressed and will not be considered in this paper.) We have recently succeeded in using CRISPR-Cas9 to create disruptive mutations in both PKA catalytic genes (PKA double KO or PKA dKO) in vasopressin-responsive kidney epithelial cells (mpkCCD cells) (9). We then used phosphoproteomics to identify a large number of novel PKA targets as well as many secondary changes in phosphorylation due to loss of PKA-mediated regulation of other kinases and phosphatases (9). Whether the two PKA catalytic proteins have redundant functions, as is implicitly assumed in many studies involving PKA-mediated regulation, has not been tested. Here, we carry out mass spectrometry-based quantitative proteomics and phosphoproteomics in both PKA catalytic-α and PKA catalytic-β single knockouts in mpkCCD collecting duct cells to address this issue. Thus, the goal of this study is to identify the relative roles of PKA-Cα and PKA-Cβ in the signaling processes triggered by vasopressin.

## METHODS

### Cell culture

The study utilized immortalized mpkCCD cells in which either *Prkaca* or *Prkacb* gene expression was deleted (“PKA-Cα-null” and “PKA-Cβ-null”) by introducing mutations using CRISPR-Cas9 (9). Control cell lines (“PKA-intact”) were carried through the CRISPR-Cas9 protocol but did not show deletion of either PKA catalytic gene. We used three independent PKA-Cα-null lines and three independent PKA-intact controls. Similarly, we used three independent PKA-Cβ-null lines and three different independent PKA-intact controls, giving 3 biological replicates for each kinase. Experiments with these lines were performed in duplicate (i.e. 2 technical replicates each) giving 24 samples in total. The overall scheme is summarized in **Figure 1**. Cells were cultured as described previously (13). Briefly, they were initially maintained in complete medium, DMEM/F-12 containing 2% serum and other supplements (5 μg/mL insulin, 50 nM dexamethasone, 1 nM triiodothyronine, 10 ng/mL epidermal growth factor, 60 nM sodium selenite, 5 μg/mL transferrin; all from Sigma). Cells were seeded on permeable membrane supports (Transwell) and grown in complete medium containing 0.1 nM 1-desamino-8-d-arginine-vasopressin (dDAVP, basal side only) for 4 days during which the cells formed confluent monolayers. Then, the medium was changed to simple medium (DMEM/F12 with dexamethasone, sodium selenite, and transferrin and no serum) with 0.1 nM dDAVP and maintained for 3 days at which time the cells were harvested for proteomic analysis. Because dDAVP was present in the culture medium under all conditions, it was not an experimental variable in the present study.

**Figure 1.**
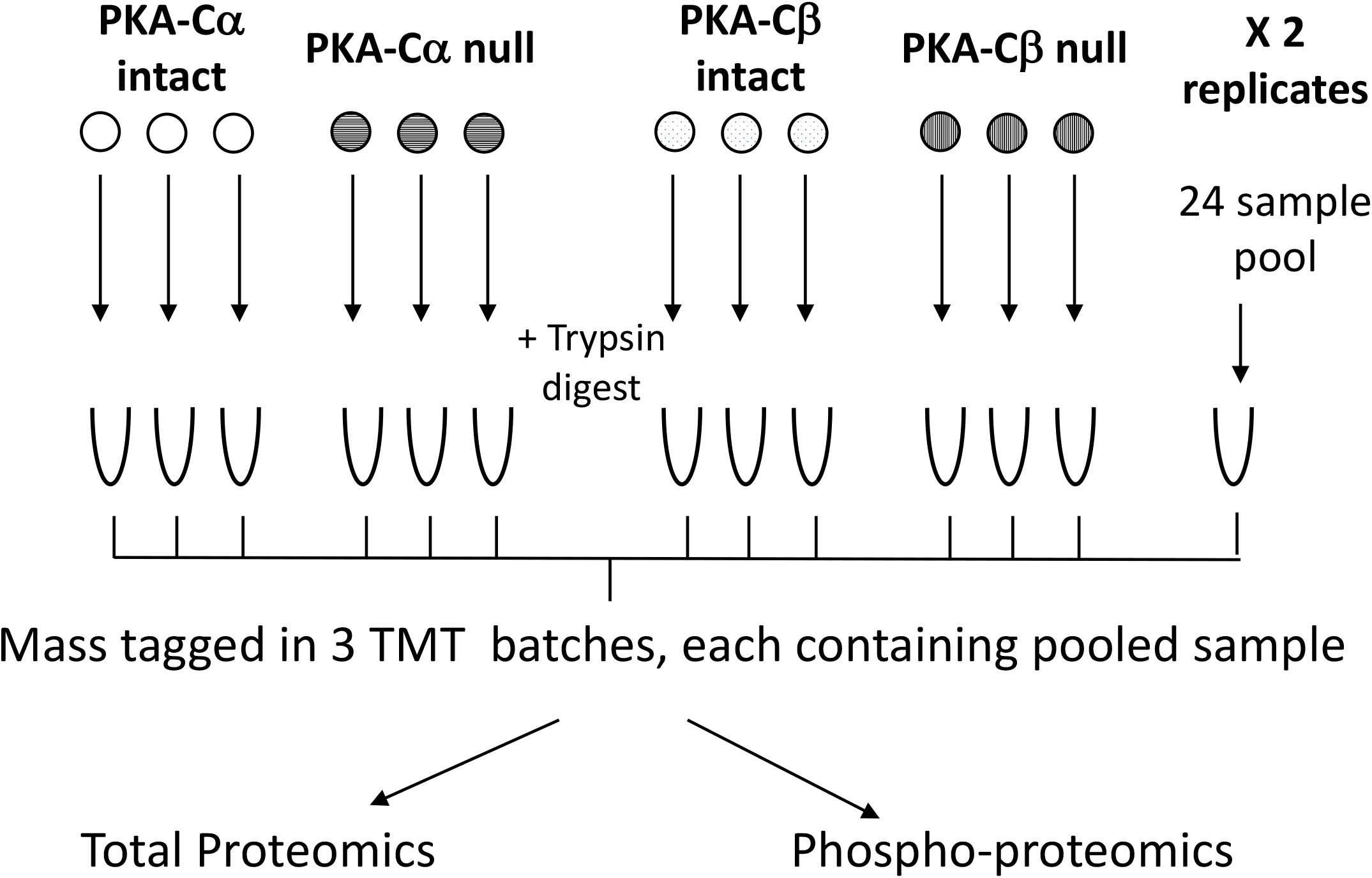
Experimental method. Cells were grown on permeable supports with three biological replicates as indicated. There were two technical replicates each resulting in a total of 24 samples. Proteins were isolated, trypsinized and labeled using isotopic tags (TMT-11 plex). The 24 samples were combined into three batches of 8 each with a shared sample pooled from all 24 samples. The combined samples were processed for proteomic and phosphoproteomic analysis as described in Methods.

### Phosphoproteomics

#### Sample preparation for total- and phospho-proteomics

Cells were washed three times with ice-cold PBS and lysed with TEAB buffer (ThermoFisher) with SDS (1%) containing protease and phosphatase inhibitors (*Halt*^*™*^, ThermoFisher). The membranes were scraped and samples were homogenized using a QIAshredder (Qiagen). Protein concentrations were measured using the Pierce™ BCA Protein Assay Kit. Protein lysates were reduced with 20 mM dithiothreitol for 1 hour at 25°C, and then alkylated with 40 mM iodoacetamide for 1 hour at 25°C in the dark. The proteins were acetone precipitated prior to digestion with recombinant trypsin (ThermoFisher) (1:50 wt/wt.) overnight at 37°C. The resulting peptides were desalted using hydrophilic-lipophilic-balanced (HLB) extraction cartridges (Oasis) and quantified using Pierce™ Quantitative Colorimetric Peptide Assay. For each replicate, 250 μg of peptide were labeled using TMT11Plex Mass Tag Labeling Kit (Thermo Scientific, Lot number UE283355) following the manufacturer’s instructions and they were combined as shown in **Figure 1**. A total of three labeling batches using the same TMT11 Plex Mass Tag kit were run to quantify all 24 samples. Each batch included a common pooled sample containing a mixture of all 24 experimental samples. Samples were combined and, after taking an aliquot for total proteomics, phosphopeptides were sequentially enriched following the Sequential Enrichment from Metal Oxide Affinity Chromatography protocol (SMOAC, Thermo Scientific). We then fractionated the samples (12 fractions) using high pH reverse phase chromatography (Agilent 1200 HPLC System). Samples were then vacuum-dried and stored at −80°C until analysis.

The dried peptides were re-suspended with 0.1% formic acid, 2% acetonitrile in LC-MS grade water (J.T. Baker) before mass spectrometry analysis. Peptides (total and phospho-) were analyzed using a Dionex UltiMate 3000 nano LC system connected to an Orbitrap Fusion Lumos mass spectrometer equipped with an EASY-Spray ion source (Thermo Fisher Scientific). Peptides were introduced into a peptide nanotrap at a flow rate of 5 μL/min. The trapped peptides were fractionated with a reversed-phase EASY-Spray PepMap column (C18, 75 μm × 50 cm) using a linear gradient of 4 to 32% acetonitrile in 0.1% formic acid (120 min at 0.3 μL/min).

#### Mass spectrometry data processing

Raw mass spectra were searched against the *Mus musculus* UniProtKB(26) reference proteome (Proteome ID: UP000000589, release 2019_06, plus contaminant database) using MaxQuant(2) 1.6.7.0. Quantitative abundance profiling of tandem mass tag (TMT)-labeled proteins and phosphopeptides were based on Synchronous Precursor Selection-MS3 (SPS-MS3) workflow (16). Lot-specific TMT isotopic impurity correction factors were used as recommended in the TMT product data sheets. Variable modifications included phospho (STY), oxidation (M), and acetyl (Protein-N-terminal). False discovery rate was controlled at 1% (target-decoy). “Trypsin/P” was set as the digestion enzyme with up to 2 missed cleavages allowed. Other parameters were set to the defaults. We used “proteinGroups.txt” output file as the input data for total proteome analyses. Both “Phospho (STY)Sites.txt” and “evidence.txt” output files were used for phosphoproteome analyses. For each TMT batch, corrected reporter ion intensities were first normalized to make the total sum intensities for each channel equal. Then, the average of the common pooled channel from three TMT batches were used to normalize the batch effects.

#### In vitro phosphorylation experiments

Three independent PKA dKO cell lines (9) were grown on permeable membrane supports as described above. The confluent monolayers were washed three times with ice-cold PBS and lysed with TEAB buffer (ThermoFisher) with SDS (1%) containing protease and phosphatase inhibitors. The membranes were scraped and samples were homogenized using a QIAshredder (Qiagen) and the proteins were precipitated with acetone. The protein pellet was resuspended in TEAB buffer and protein concentration was determined by Pierce™ BCA Protein Assay Kit.

Equal quantities of proteins from the three PKA dKO cell lines were pooled together and 500 μg pooled protein extract was mixed with either recombinant PKA-Cα (1.5μg) or recombinant PKA-Cβ (1.5μg) enzymes obtained from Genetex Inc. (PRKACA-GTX65206-pro; PRKACB-GTX65207-pro). Samples were incubated at 30° C for 24 h in a buffer containing 50 mM Tris-HCl, 10 mM MgCl_2,_ 0.1 mM EDTA, 2 mM DTT and 250 μM ATP. Samples without any added kinases were used as controls. Following the in vitro kinase incubation, the proteins underwent a standard phosphoproteomic analysis as described previously.

#### Bioinformatics and data sharing

Technical replicates were summarized by taking the median value. All analyses were performed using Perseus, Excel, and R software. Moderated P-values (P_mod_) were calculated using the empirical Bayes method, which integrates variance information from all peptides measured in the same LC-MS/MS run (10). Data have been deposited in PRIDE (as part of ProteomeXchange) with accession number PXD015050 (URL for reviewer is https://www.ebi.ac.uk/pride/archive/projects/PXD015050.)

## RESULTS

Previously, the PKA-Cα-null and PKA-Cβ-null cells were characterized by immunoblotting, which showed that PKA-Cα and PKA-Cβ proteins were undetectable in the corresponding single knockout lines (9). Both single KO lines grew well and formed confluent mono-layered epithelial sheets when grown on permeable supports. Here, we used protein mass spectrometry to quantify changes in total protein abundances and changes in phosphorylation in PKA-Cα-null versus -intact, and PKA-Cβ-null versus - intact cells across the proteome.

### Proteome-wide quantification of protein abundances

Quantitative data for the total abundance of all individual proteins is provided on a specially designed web page (https://hpcwebapps.cit.nih.gov/ESBL/Database/PKA-singleKO-total/) and as **Supplemental Dataset 1,** provided at https://hpcwebapps.cit.nih.gov/ESBL/Database/PKA-singleKO-total/. **Figure 2** shows changes in protein abundances corresponding to 4691 different genes in both PKA-Cα-null cells and PKA-Cβ-null cells versus their respective PKA-intact control cells. Each point shows values for a different protein. The proteins whose abundances were previously found to be significantly altered in PKA-Cα/PKA-Cβ double knockout (dKO) cells are marked by red or green points depending on the direction of change in the dKO. Interestingly, aquaporin-2 (Aqp2), which had previously been seen to be almost completely ablated in the PKA dKO cells, was decreased in only the PKA-Cα-null cells and not the PKA-Cβ-null cells. Two other proteins that were strongly decreased in abundance in the PKA dKO experiments, namely Complement Factor C3 (C3) and Mucin-4 (Muc4) were differentially affected in the two single KO cells. C3 was selectively decreased in PKA-Cα-null cells, whereas Mucin-4 was selectively decreased in PKA-Cβ-null cells. These data support the view that PKA-Cα and PKA-Cβ do not have the same physiological roles in collecting duct cells. Interestingly, PKA-Cβ was substantially increased in abundance in the PKA-Cα-null cells suggesting a compensatory response.

**Figure 2.**
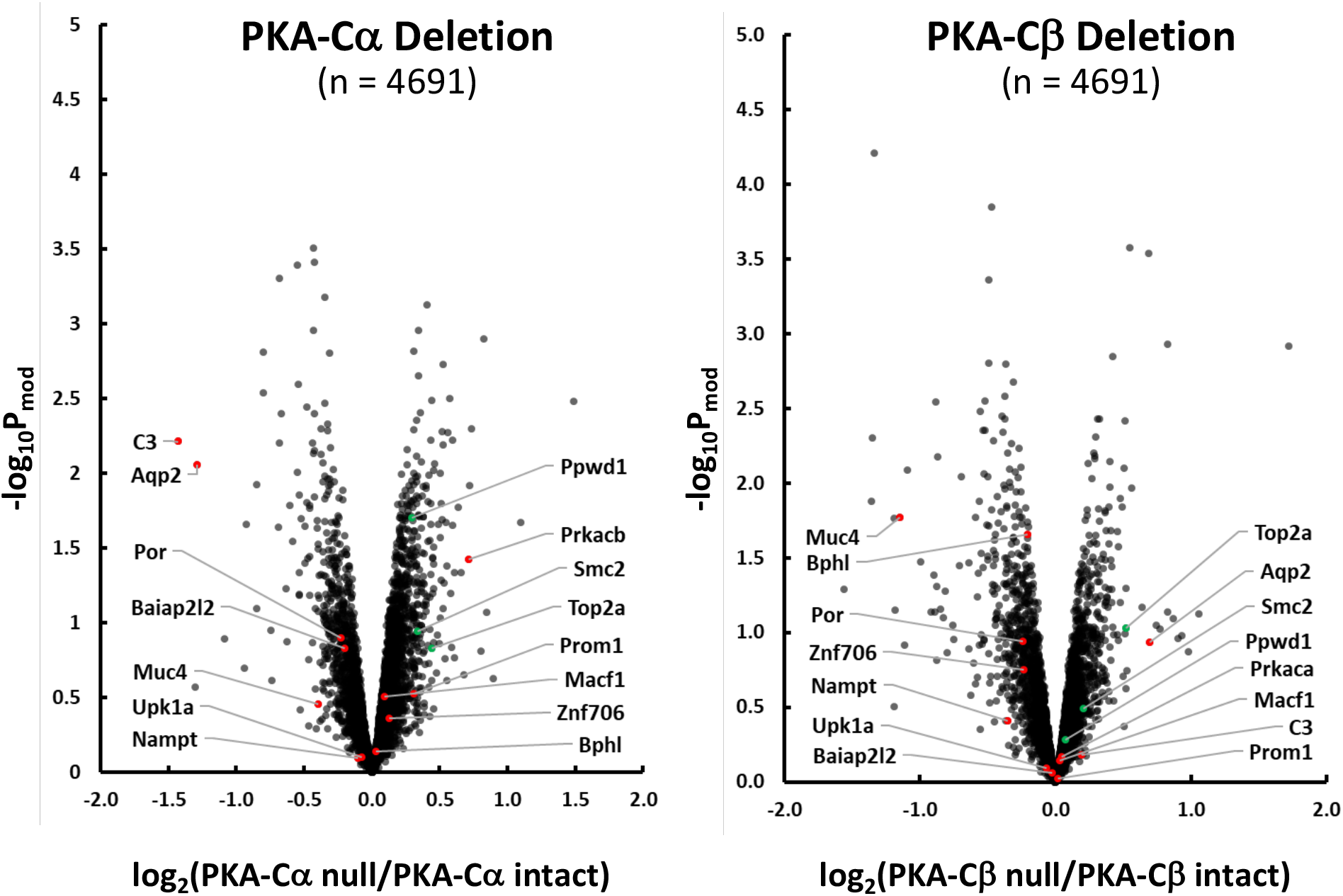
Effect of PKA-Cα (left) and PKA-Cβ (right) deletion on protein abundances in mouse mpkCCD cells. Red points indicate proteins decreased in PKA-Cα/PKA-Cβ double knockout cells (9). Green points indicate proteins increased in PKA-Cα/PKA-Cβ double knockout cells (9). Arrows to these red and green points show official gene symbols for the specific proteins.

We plotted the changes in protein abundances in the PKA-Cα-null cells versus those in the PKA-Cβ-null cells for those proteins that changed significantly in either PKA-KO line (**Figure 3A**). As can be seen, some proteins (yellow) underwent parallel changes in abundance in both single KO cell lines, while others underwent selective, significant changes in either PKA-Cα-null (red) or PKA-Cβ-null (green) cells. Control vs. control comparisons for the same proteins are shown in **Figure 3B** illustrating the magnitude of changes expected from random variability in the mass spectrometric quantification. (The 95% confidence interval for log_2_(control/control) was 0.382). A listing of the proteins that most convincingly changed (P_mod_ < 0.02 and |log(ratio)| > 0.5) in the PKA-Cα-null cells is given in **Table 1**, while changes in PKA-Cβ-null cells are listed in **Table 2.** Overall, we conclude that PKA-Cα and PKA-Cβ are not redundant with respect to regulation of protein abundances.

**Table 1.** Proteins with substantial changes in abundance in response to PKA-Cα deletion (P_mod_ < 0.02 and |log(ratio)| > 0.5).

| UniProt ID | Gene Symbol | Annotation | Log<sub>2</sub><br>(PKA-Cα<br>null/ PKA-<br>Cα intact) | Pmod | Log<sub>2</sub><br>(PKA-Cβ<br>null/ PKA-<br>Cβ intact) | Pmod | Log<sub>2</sub><br>(PKA-Cα<br>null/ PKA-<br>Cβ null) |
| --- | --- | --- | --- | --- | --- | --- | --- |
| P01027 | **C3** | Complement C3 | **-1.427** | 0.006 | 0.195 | 0.655 | -0.990 |
| P56402 | **Aqp2** | Aquaporin-2 | **-1.286** | 0.009 | 0.694 | 0.115 | -1.520 |
| Q9QYI5 | **Dnajb2** | DnaJ homolog subfamily B member 2 | **-0.850** | 0.012 | 0.056 | 0.848 | -0.513 |
| P97742 | **Cpt1a** | Carnitine O-palmitoyltransferase 1, liver isoform | **-0.799** | 0.002 | -0.091 | 0.647 | -0.467 |
| Q8BH86 | **Dglucy** | D-glutamate cyclase, mitochondrial | **-0.798** | 0.003 | -0.094 | 0.666 | -0.090 |
| P24472 | **Gsta4** | Glutathione S-transferase A4 | **-0.684** | 0.006 | -0.451 | 0.049 | 0.199 |
| Q9EPK8 | **Trpv4** | Transient receptor potential channel subfamily V member 4 | **-0.682** | 4.93E-04 | 0.067 | 0.647 | -0.435 |
| Q9WTQ5 | **Akap12** | A-kinase anchor protein 12 | **-0.669** | 0.004 | 0.070 | 0.714 | -0.379 |
| Q99J39 | **Mlycd** | Malonyl-CoA decarboxylase, mitochondrial | **-0.602** | 0.016 | 0.087 | 0.690 | -0.245 |
| Q8BLF1 | **Nceh1** | Neutral cholesterol ester hydrolase 1 | **-0.551** | 0.010 | -0.346 | 0.077 | 0.072 |
| P63011 | **Rab3a** | Ras-related protein Rab-3A | **-0.550** | 4.05E-04 | -0.115 | 0.324 | -0.186 |
| P26443 | **Glud1** | Glutamate dehydrogenase 1, mitochondrial | **-0.543** | 0.003 | -0.057 | 0.692 | -0.376 |
| G3X9Y6 | **Akr1c19** | Aldo-keto reductase family 1, member C19 | **-0.534** | 0.014 | -0.307 | 0.123 | -0.086 |
| Q8BG05 | **Hnrnpa3** | Heterogeneous nuclear ribonucleoprotein A3 | **0.507** | 0.010 | 0.033 | 0.845 | 0.162 |
| D3YYU8 | **Obsl1** | Obscurin-like protein 1 | **0.512** | 0.016 | 0.024 | 0.897 | 0.332 |
| P62482 | **Kcnab2** | Voltage-gated potassium channel subunit beta-2 | **0.518** | 0.005 | 0.366 | 0.033 | 0.062 |
| Q8VI84 | **Noc3l** | Nucleolar complex protein 3 homolog | **0.524** | 0.002 | 0.369 | 0.016 | 0.029 |
| Q5DU09 | **Znf652** | Zinc finger protein 652 | **0.527** | 0.006 | -0.275 | 0.108 | 0.396 |
| O88508 | **Dnmt3a** | DNA (cytosine-5)-methyltransferase 3A | **0.562** | 0.005 | -0.034 | 0.838 | 0.198 |
| O54786 | **Dffa** | DNA fragmentation factor subunit alpha | **0.576** | 0.003 | 0.255 | 0.127 | 0.270 |
| Q9D187 | **Ciao2b** | Cytosolic iron-sulfur assembly component 2B | **0.596** | 0.006 | -0.206 | 0.268 | 0.422 |
| Q61180 | **Scnn1a** | Amiloride-sensitive sodium channel subunit alpha | **0.637** | 0.017 | -0.016 | 0.945 | 0.616 |
| P0DOV2 | **Ifi204** | Interferon-activable protein 204 | **0.722** | 0.012 | 0.108 | 0.664 | 0.224 |
| Q9JK42 | **Pdk2** | Pyruvate dehydrogenase kinase isozyme 2 | **0.734** | 0.005 | 0.049 | 0.821 | 0.167 |
| P26645 | **Marcks** | Myristoylated alanine-rich C-kinase substrate | **1.486** | 0.003 | 1.718 | 0.001 | 0.089 |
| P05132 | **Prkaca** | cAMP-dependent protein kinase catalytic subunit $\alpha$ | - | - | 0.049 | 0.677 | - |

**Table 2.** Proteins with substantial changes in abundance in response to PKA-Cβ deletion (P_mod_ < 0.02 and |log(ratio)| > 0.5).

| UniProt ID | Gene Symbol | Annotation | Log<sub>2</sub><br>(PKA-Cα<br>null/ PKA-<br>Cα intact) | Pmod | Log<sub>2</sub><br>(PKA-Cβ<br>null/ PKA-<br>Cβ intact) | Pmod | Log<sub>2</sub><br>(PKA-Cα<br>null/ PKA-<br>Cβ null) |
| --- | --- | --- | --- | --- | --- | --- | --- |
| Q8VCT4 | **Ces1d** | Carboxylesterase 1D | -0.370 | 0.440 | **-1.358** | 0.013 | 0.731 |
| Q8CEZ4 | **Mab21l4** | Protein mab-21-like 4 | 0.180 | 0.652 | **-1.351** | 0.005 | 1.130 |
| P30115 | **Gsta3** | Glutathione S-transferase A3 | -0.294 | 0.202 | **-1.334** | 6.23E-05 | 1.129 |
| P08074 | **Cbr2** | Carbonyl reductase [NADPH] 2 | -0.178 | 0.683 | **-1.186** | 0.017 | 0.686 |
| Q8JZM8 | **Muc4** | Mucin-4 | -0.396 | 0.353 | **-1.144** | 0.017 | 0.696 |
| P00329 | **Adh1** | Alcohol dehydrogenase 1 | 0.334 | 0.348 | **-1.094** | 0.008 | 1.375 |
| Q7TPW1 | **Nexn** | Nexilin | 0.228 | 0.349 | **-0.881** | 0.003 | 0.724 |
| Q9D939 | **Sult1c2** | Sulfotransferase 1C2 | -0.535 | 0.065 | **-0.869** | 0.007 | 0.643 |
| Q62469 | **Itga2** | Integrin alpha-2 | -0.074 | 0.743 | **-0.696** | 0.009 | 0.684 |
| Q99LB7 | **Sardh** | Sarcosine dehydrogenase, mitochondrial | -0.163 | 0.457 | **-0.591** | 0.017 | 0.530 |
| Q9WUU7 | **Ctsz** | Cathepsin Z | -0.053 | 0.786 | **-0.562** | 0.013 | 0.585 |
| Q8R3G9 | **Tspan8** | Tetraspanin-8 | -0.199 | 0.212 | **-0.557** | 0.003 | 0.446 |
| P22935 | **Crabp2** | Cellular retinoic acid-binding protein 2 | -0.282 | 0.088 | **-0.534** | 0.004 | 0.120 |
| P63082 | **Atp6v0c** | V-type proton ATPase 16 kDa proteolipid subunit | -0.101 | 0.556 | **-0.526** | 0.009 | 0.401 |
| Q80V42 | **Cpm** | Carboxypeptidase M | -0.077 | 0.607 | **-0.516** | 0.004 | 0.408 |
| P12265 | **Gusb** | Beta-glucuronidase | -0.032 | 0.819 | **-0.516** | 0.003 | 0.496 |
| P17563 | **Selenbp1** | Methanethiol oxidase | -0.122 | 0.525 | **-0.506** | 0.019 | 0.228 |
| Q5FWI3 | **Cemip2** | Cell surface hyaluronidase | 0.020 | 0.899 | **0.508** | 0.008 | 0.281 |
| P55264 | **Adk** | Adenosine kinase | -0.297 | 0.059 | **0.513** | 0.004 | -0.689 |
| Q99PT3 | **Ino80b** | INO80 complex subunit B | 0.048 | 0.658 | **0.551** | 2.66E-04 | -0.279 |
| Q8K0C4 | **Cyp51a1** | Lanosterol 14-alpha demethylase | -0.313 | 0.117 | **0.563** | 0.011 | -0.479 |
| P28661 | **Septin4** | Septin-4 | 0.235 | 0.106 | **0.690** | 2.90E-04 | -0.229 |
| Q80Z25 | **Ofd1** | Oral-facial-digital syndrome 1 protein homolog | 0.518 | 0.020 | **0.826** | 0.001 | -0.008 |
| P26645 | **Marcks** | Myristoylated alanine-rich C-kinase substrate | 1.486 | 0.003 | **1.718** | 0.001 | 0.089 |
| P68181 | **Prkacb** | cAMP-dependent protein kinase catalytic subunit beta | 0.717 | 0.038 | - | - | - |

**Figure 3.**
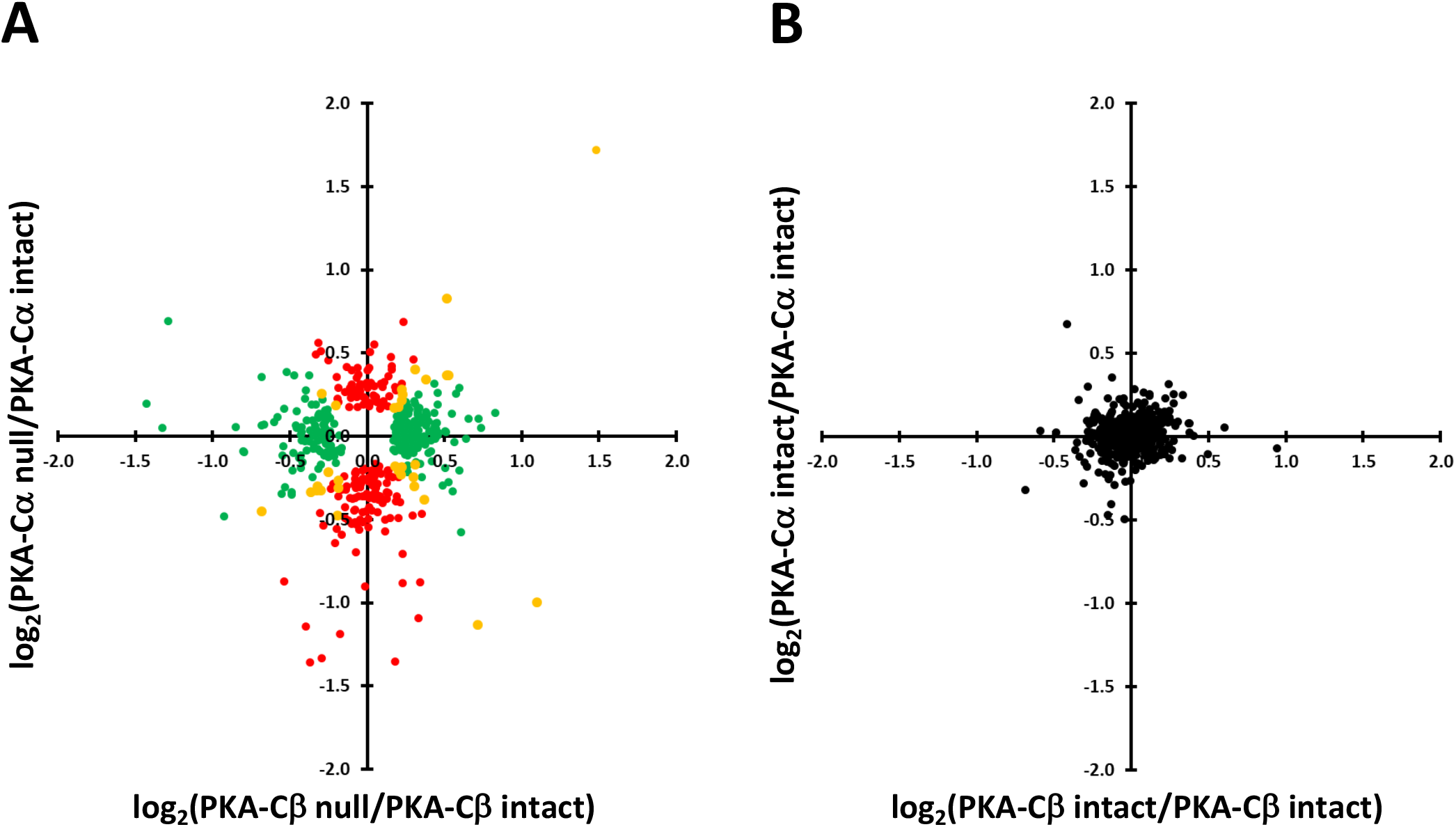
Comparison of effects of PKA-Cα and PKA-Cβ deletion on protein abundances. A. Red points show proteins changed with PKA-Cα deletion, but not PKA-Cβ deletion. Green points show proteins changed with PKA-Cβ deletion, but not PKA-Cα deletion. Yellow points are changed in both, but not necessarily in the same direction. B. Intrinsic variability of the data estimated by comparing values for PKA-intact controls versus other controls (see text).

### Proteome-wide quantification of protein phosphorylation

Quantitative data for all phosphopeptides are provided at (https://hpcwebapps.cit.nih.gov/ESBL/Database/PKA-singleKO-phospho/) and as **Supplemental Dataset 2**. **Figure 4** shows changes in phosphopeptide abundances (n= 4635) in both PKA-Cα-null cells and PKA-Cβ-null cells versus their respective PKA-intact (control) cells. Each point shows values for a different phosphopeptide. The phosphopeptides corresponding to previously identified PKA target sites (9) are indicated in red. Most of the PKA sites were found to be decreased in PKA-Cα-null but not in PKA-Cβ-null cells relative to controls. A listing of the phosphopeptides most convincingly changed (P_mod_ < 0.01 and |log(ratio)| > 0.5) in the PKA-Cα-null cells is given in **Table 3**, while changes in PKA-Cβ-null cells are listed in **Table 4.** Overall, we conclude that PKA-Cα and PKA-Cβ have a substantially different set of phosphorylation targets. This difference could be due either to a difference in target sequence preference or a difference in PKA-Cα and PKA-Cβ interactomes. **Figure 5** shows the predicted target sequence preference logos for decreased single-site phosphopeptides in PKA-Cα- and PKA-Cβ-null cells, respectively. Only the PKA-Cα logo was consistent with the familiar (R/K)-(R/K)-X-p(S/T) motif generally accepted to be characteristic of PKA. Among the 82 phosphosites that were decreased in the PKA-Cβ-null cells, only four possessed the (R/K)-(R/K)-X-p(S/T) motif. The logos raise the hypothesis that the two PKA catalytic subunits may have different target sequence preferences.

**Table 3.**
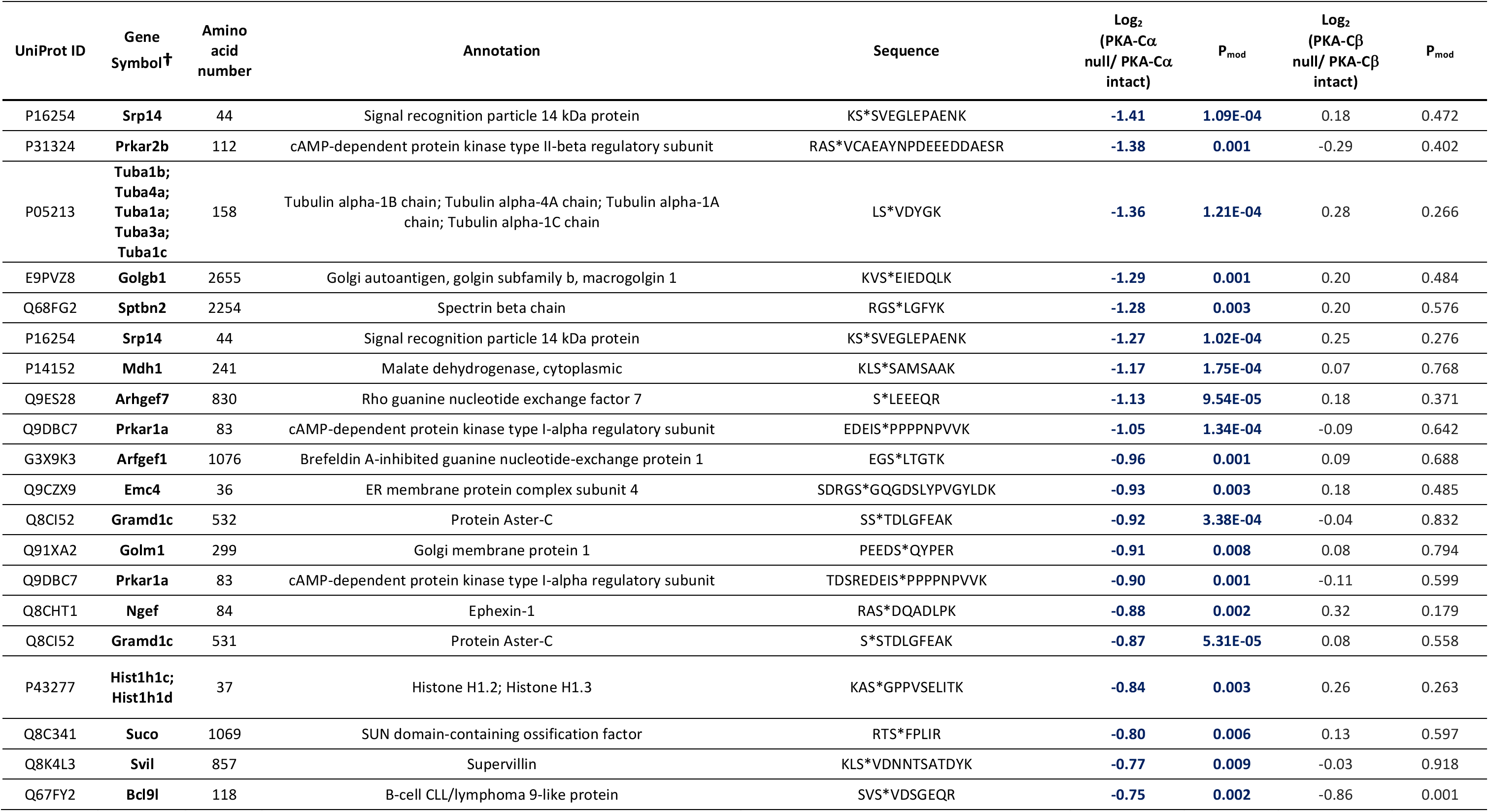
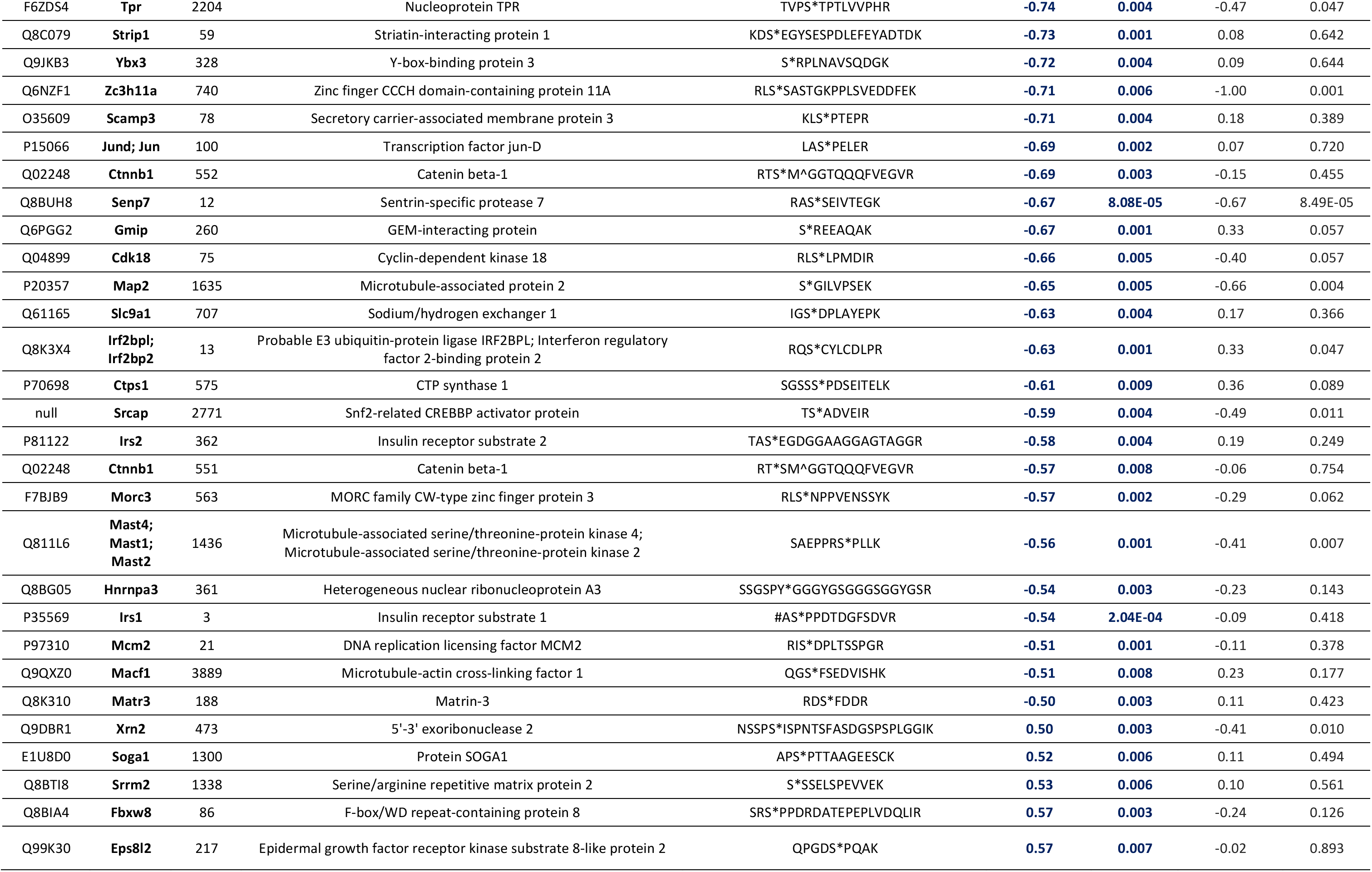
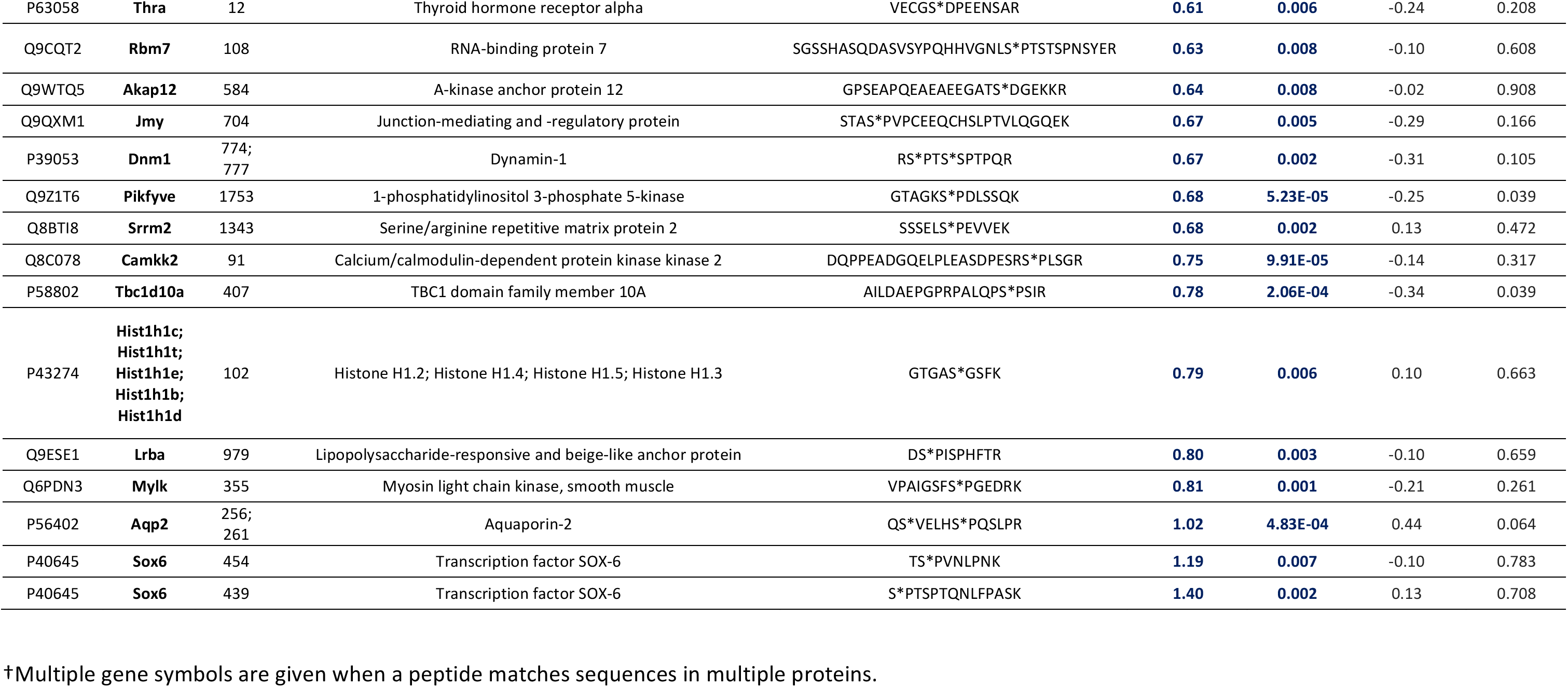
Phosphopeptides most convincingly changed in PKA-Cα-null cells versus PKA-Cα-intact (P_mod_ < 0.01 and |log(ratio)| > 0.5). PKA-Cβ-null data for the same peptides is also given for comparison.

**Table 4.** Phosphopeptides changed most convincingly in PKA-Cβ null cells versus PKA-Cβ intact (P_mod_ < 0.01 and |log(ratio)| > 0.5). PKA-Cα-null data for the same peptides is also given for comparison.

| UniProt ID | Gene Symbol | Amino acid number | Annotation | Sequence | Log2 (PKA-Cα null/ PKA-Cα intact) | $P_{\text{mod}}$ | Log2 (PKA-Cβ null/ PKA-Cβ intact) | $P_{\text{mod}}$ |
| --- | --- | --- | --- | --- | --- | --- | --- | --- |
| Q6NZF1 | **Zc3h11a** | 740 | Zinc finger CCCH domain-containing protein 11A | RLS*SASTGKPPLSVEDDFEK | -0.71 | 0.006 | **-1.00** | **0.001** |
| Q67FY2 | **Bcl9l** | 118 | B-cell CLL/lymphoma 9-like protein | SVS*VDSGEQR | -0.75 | 0.002 | **-0.86** | **0.001** |
| Q03173 | **Enah** | 719 | Protein enabled homolog | APST*STPEPTR | 0.00 | 0.988 | **-0.79** | **0.002** |
| Q9QZQ1 | **Mllt4** | 1201 | Afadin | ITSVS*TGNLCTEEQSPPRPEAYPIPTQTYTR | -0.05 | 0.793 | **-0.77** | **0.002** |
| O08796 | **Eef2k** | 66 | Eukaryotic elongation factor 2 kinase | T*ECGSTGSPASSFHFK | -0.13 | 0.552 | **-0.75** | **0.004** |
| Q03173 | **Enah** | 720 | Protein enabled homolog | APSTS*TPEPTR | -0.11 | 0.633 | **-0.75** | **0.005** |
| O55003 | **Bnip3** | 60 | BCL2/adenovirus E1B 19 kDa protein-interacting protein 3 | SSHCDs*PPR | 0.27 | 0.222 | **-0.70** | **0.007** |
| Q2VPU4 | **MLxip** | 9 | MLX-interacting protein | #AADVFM^CS*PR | -0.03 | 0.897 | **-0.68** | **0.010** |
| Q8BUH8 | **Senp7** | 12 | Sentrin-specific protease 7 | RAS*SEIVTEGK | -0.67 | 8.08E-05 | **-0.67** | **8.49E-05** |
| P20357 | **Map2** | 1635 | Microtubule-associated protein 2 | S*GILVPSEK | -0.65 | 0.005 | **-0.66** | **0.004** |
| Q8VDZ4 | **Zdhhc5** | 409 | Palmitoyltransferase ZDHHC5 | SEPSLEPESFRS*PTFGK | -0.27 | 0.144 | **-0.64** | **0.003** |
| Q8CGF1 | **Arhgap29** | 1241 | Rho GTPase-activating protein 29 | ESSEEPALPEGT*PTCQRPR | -0.04 | 0.743 | **-0.63** | **0.001** |
| Q8VI36 | **Pxn** | 258 | Paxillin | IS*ASSATR | -0.01 | 0.966 | **-0.62** | **0.004** |
| Q8C6B2 | **Rtkn** | 518 | Rhotekin | T*FSLDAAPADHSLGPSR | -0.01 | 0.962 | **-0.60** | **0.005** |
| Q8C6B2 | **Rtkn** | 520 | Rhotekin | TFS*LDAAPADHSLGPSR | -0.13 | 0.423 | **-0.60** | **0.002** |
| A1A535 | **Veph1** | 422 | Ventricular zone-expressed PH domain-containing protein 1 | INAESNT*PGSGR | -0.34 | 0.070 | **-0.59** | **0.005** |
| Q9Z2H5 | **Epb41l1** | 540 | Band 4.1-like protein 1 | RLPS*SPASPSPK | -0.17 | 0.332 | **-0.57** | **0.006** |
| P28661 | **Sept4** | 68 | Septin-4 | PQS*PDLCDDVEFR | -0.02 | 0.917 | **-0.55** | **0.003** |
| Q8C5R2 | **Proser2** | 223 | Proline and serine-rich protein 2 | LAGNEALSPTS*PSK | -0.26 | 0.119 | **-0.55** | **0.004** |
| Q99MR1 | **Gigyf1** | 227 | GRB10-interacting GYF protein 1 | ST*SPDGGPR | -0.05 | 0.640 | **-0.51** | **0.001** |
| Q99MR1 | **Gigyf1** | 226 | GRB10-interacting GYF protein 1 | S*TSPDGGPR | 0.05 | 0.723 | **-0.50** | **0.003** |
| Q60864 | **Stip1** | 481 | Stress-induced-phosphoprotein 1 | HDS*PEDVK | -0.12 | 0.498 | **0.57** | **0.005** |

**Figure 4.**
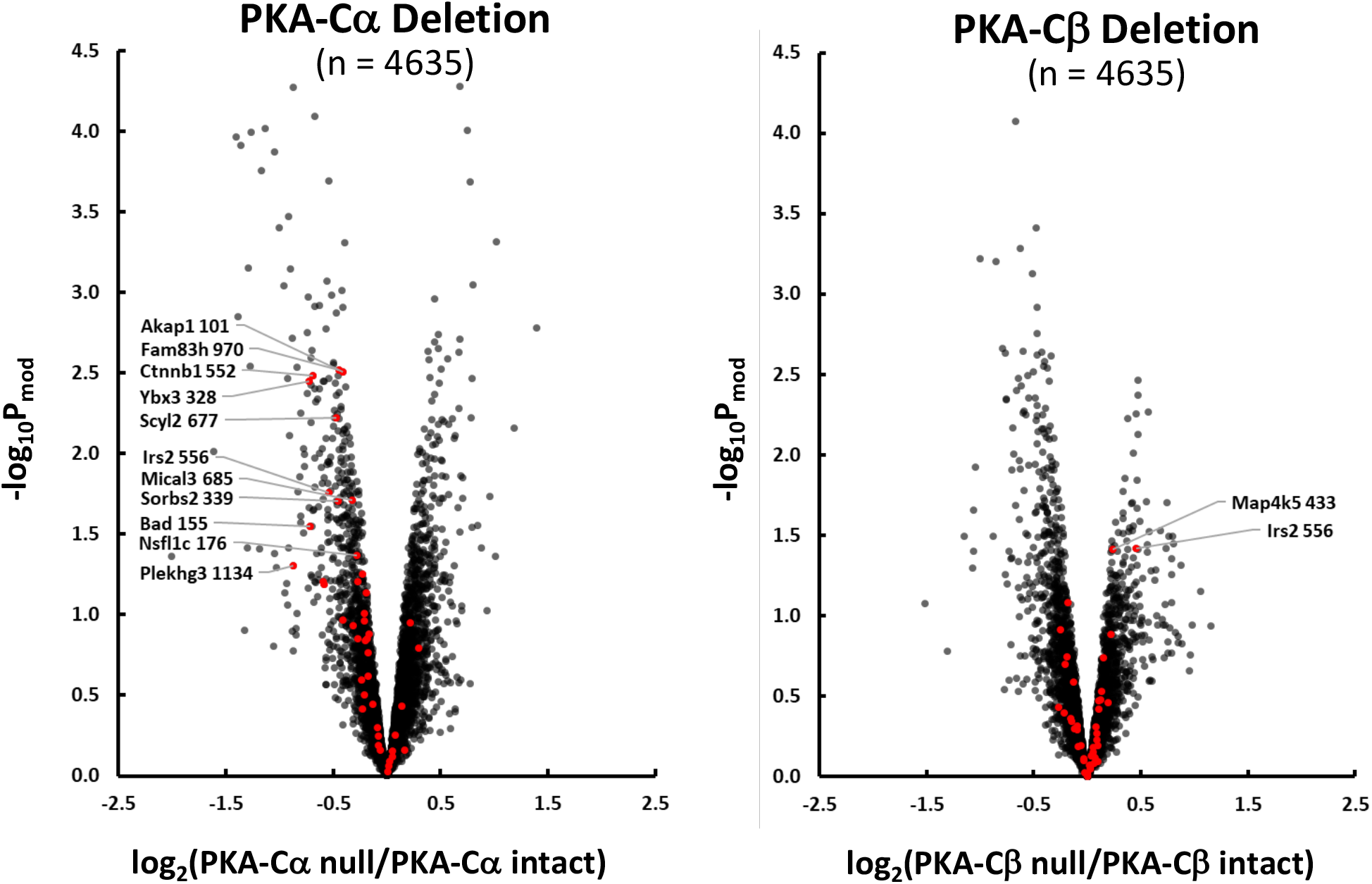
Effect of PKA-Cα (left) and PKA-Cβ (right) deletion on phosphopeptide abundances in mouse mpkCCD cells. Red points indicate phosphorylation sites altered in PKA-Cα/PKA-Cβ double knockout cells (9). Arrows to these red points show official gene symbol and amino acid number of phosphorylated site for those sites above P_mod_ threshold (−log P_mod_ > 1.3).

**Figure 5.**
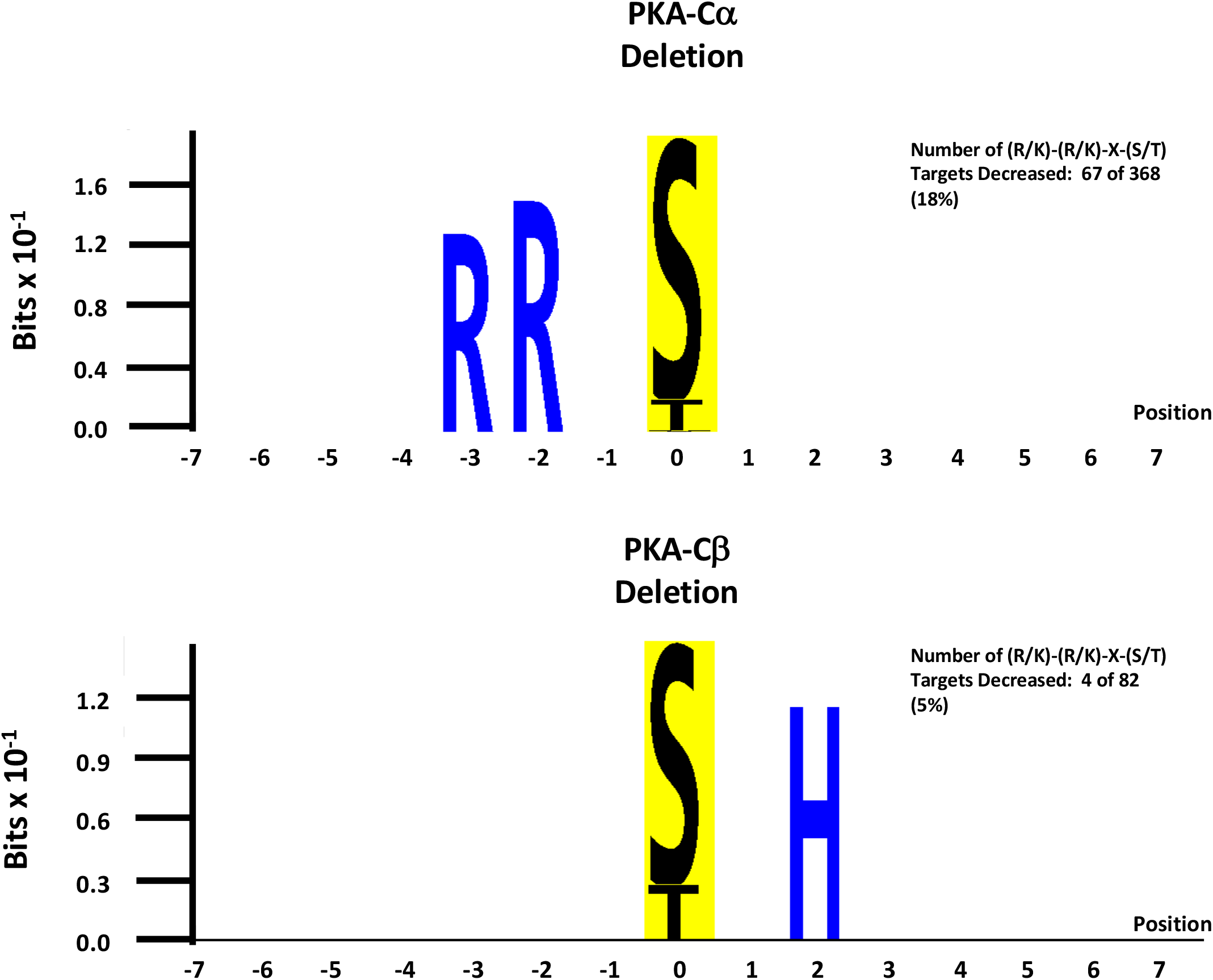
Sequence preference logos from sites decreased with PKA-Cα deletion (top) and PKA-Cβ deletion (bottom). Logos were generated using *PTM-Logo* with a background of all unchanged phosphopeptides.

### In vitro phosphorylation by recombinant PKA-Cα versus PKA-Cβ

PKA-Cα and PKA-Cβ may have different target preferences. To test this hypothesis directly, we carried out in-vitro phosphorylation studies using purified recombinant PKA-Cα and PKA-Cβ catalytic subunits to phosphorylate protein extracts obtained from PKA-dKO cells. Full data are in **Supplemental Dataset 3. Figure 6A** comparing the changes in phosphorylation by PKA-Cα versus PKA-Cβ shows that both PKA catalytic subunits phosphorylate virtually identical substrates. **Figure 6B** shows that the sequence preferences for PKA-Cα versus PKA-Cβ are virtually identical. This result rules out the hypothesis that the two PKA catalytic subunits have different substrate specificities. The remaining possibility, that the two catalytic subunits have different targets because they are propinquitous to a different set of proteins, remains as the most likely explanation for the differences in targets in the intact cells.

**Figure 6.**
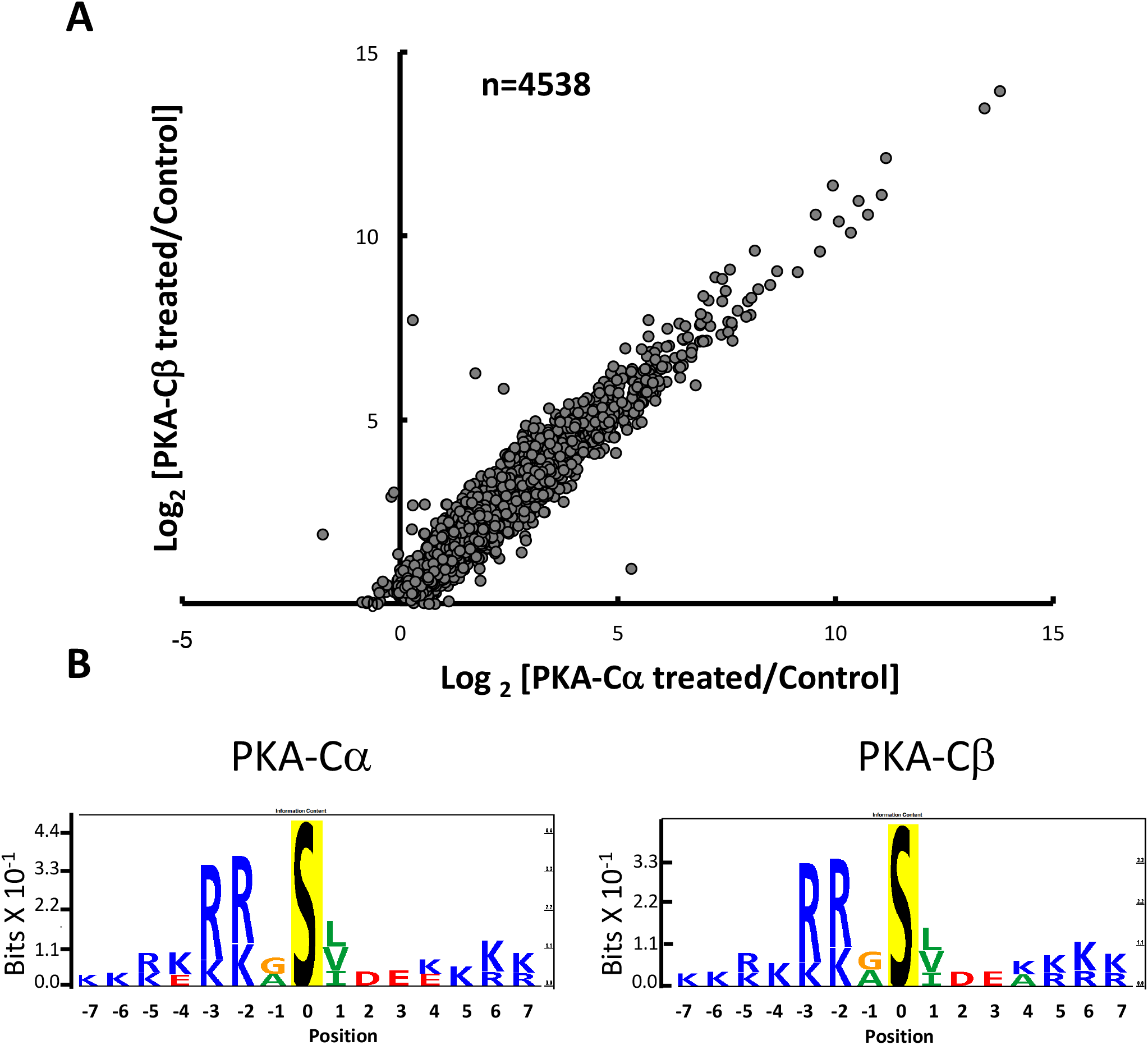
Results of *in vitro* phosphorylation experiments. Protein extracts from PKA double knockout cells (PKA-Cα and PKA-Cβ) were incubated with recombinant PKA-Cα or PKA-Cβ and phosphorylation was quantified by mass spectrometry. A. There was a marked similarity between responses to the two recombinant kinase proteins. B. Sequence preference logos derived from the analysis were almost identical.

### Phosphoproteomics as a virtual proximity assay

In addition to the target sequence, another factor important to the determination of targets for a protein kinase is co-localization because a kinase can only phosphorylate a protein with which it comes into physical contact. In this sense, phosphoproteomic analysis can be viewed as a kind of large-scale proximity assay, identifying kinase/target interactions. To evaluate whether differences in PKA-Cα and PKA-Cβ phosphorylation targets result in part from differences in PKA-Cα and PKA-Cβ localization, we identified *Gene Ontology (GO-CC) Cellular Component* terms that are enriched in either set of phosphorylation targets in the intact cell experiments, relative to all proteins detected. **Table 5** shows terms enriched in phosphoproteins with altered phosphorylation in the PKA-Cα-null cells, while **Table 6** shows terms enriched in phosphoproteins with altered phosphorylation in the PKA-Cβ-null cells. Enriched terms in the PKA-Cα-null cells were largely related to cell membranes and membrane vesicles, while enriched terms in the PKA-Cβ-null cells were related to the actin cytoskeleton and cell junctions, suggesting that in vivo cellular subdomains of the PKA-Cα and PKA-Cβ differ.

**Table 5.** *Gene Ontology Cellular Component* terms enriched in list of phosphoproteins with altered phosphorylation in PKA-Cα-null cells.

| Term | Number | Fold Enrichment | Fisher Exact | Proteins |
| --- | --- | --- | --- | --- |
| <u>membrane raft</u> | 8 | 4.3 | 3.20E-04 | EGFR, EFHD2, PRKAR2B, PRKAR1A, PIKFYVE, CXADR, CTNNB1, HDAC6 |
| <u>perinuclear region of cytoplasm</u> | 12 | 2.3 | 3.90E-03 | EGFR, PACS1, EPN3, PRKAR2B, FBXW8, PAK2, SCYL2, PIKFYVE, SPTBN2, RAB3IP, CTNNB1, HDAC6 |
| <u>membrane part</u> | 26 | 1.6 | 6.40E-03 | GPRC5C, ACSS2, CXADR, CHCHD6, CTNNB1, EFHD2, PRKAR2B, PIKFYVE, PRKAA1, SYNPO, SREBF1, EGFR, EPN3, OLF120, KANSL1, SLC33A1, STIM2, CRB3, EMC4, BNIP3L, PRKAR1A, SPTBN2, GRAMD1C, DNM1, HDAC6, SUCO |
| <u>cytoplasmic vesicle</u> | 11 | 2 | 1.90E-02 | EGFR, PACS1, SREBF1, ARHGAP21, EPN3, SCYL2, RAN, PIKFYVE, SPTBN2, CXADR, DNM1 |
| <u>integral component of membrane</u> | 16 | 1.6 | 3.60E-02 | SREBF1, EGFR, OLF120, GPRC5C, KANSL1, SLC33A1, STIM2, CRB3, ACSS2, CXADR, CHCHD6, EMC4, BNIP3L, PIKFYVE, GRAMD1C, SUCO |
| <u>membrane-bounded vesicle</u> | 21 | 1.4 | 5.00E-02 | SREBF1, PACS1, EGFR, SRP14, EPN3, GPRC5C, RAN, CRB3, CXADR, HNRNPA1, CTNNB1, ARHGAP21, FAM65A, PRKAR2B, RPL30, SCYL2, PIKFYVE, SPTBN2, TUBA1B, DNM1, MDH1 |

**Table 6.** *Gene Ontology Cellular Component* terms enriched in list of phosphoproteins with altered phosphorylation in PKA-Cβ-null cells.

| Term | Number | Fold Enrichment | Fisher Exact | Proteins |
| --- | --- | --- | --- | --- |
| <u>cytoskeleton</u> | 23 | 1.7 | 2.70E-03 | DPF2, SEPT4, ENAH, DYNC1LI2, PDLIM5, AKAP12, IGF2BP2, NEXN, MYO9A, PXN, SMC3, TNKS1BP1, EPB41L1, MACF1, MAP2, MAP4, RANBP2, CDC42EP4, CEP170B, UBXN6, AFAP1, SYNPO, SEPT9 |
| <u>adherens junction</u> | 16 | 1.9 | 5.10E-03 | EGFR, ENAH, PDLIM5, AKAP12, NEXN, CXADR, PXN, TNKS1BP1, EPB41L1, LIMD1, EEF1D, ERC1, EPS8L2, AFAP1, TJP2, SEPT9 |
| <u>cell junction</u> | 19 | 1.7 | 1.30E-02 | CLDN8, EGFR, ENAH, PDLIM5, AKAP12, CXADR, NEXN, PXN, TNKS1BP1, EPB41L1, LIMD1, EEF1D, ERC1, CDC42EP4, EPS8L2, AFAP1, TJP2, SYNPO, SEPT9 |
| <u>plasma membrane</u> | 21 | 1.5 | 1.90E-02 | CLDN8, EGFR, ENAH, IRS2, ZDHHC5, PDLIM5, LRBA, AKAP12, WWC1, ZBTB16, CXADR, PXN, AQP2, TNKS1BP1, EPB41L1, MACF1, MAP4, CDC42EP4, EPS8L2, TJP2, SYNPO |
| <u>actin cytoskeleton</u> | 9 | 2.1 | 2.60E-02 | ENAH, MACF1, PDLIM5, CDC42EP4, MYO9A, AFAP1, PXN, SEPT9, SYNPO |

### PKA-Cα and PKA-Cβ regulate different protein kinases

Many phosphorylation sites that changed in the PKA-Cα-null or PKA-Cβ-null cells did not conform to the classic PKA target motif and presumably undergo changes in phosphorylation as a result of secondary effects on other protein kinases. **Table 7** shows protein kinase catalytic proteins that underwent changes in phosphorylation in either the PKA-Cα- or PKA-Cβ-null cells. Many of the altered phosphorylation sites have known effects on enzyme activity as indicated in the last column. The affected kinases, namely calcium/calmodulin-dependent protein kinase kinase 2 (Camkk2), the epidermal growth factor receptor kinase (Egfr), myosin light chain kinase (Mylk), p21-activated kinase (Pak2), AMP-activated protein kinase α-1 (Prkaa1), and salt-inducible kinase 2 (Sik2), all underwent changes in phosphorylation in response to PKA-Cα deletion, while only Egfr underwent a change in phosphorylation in response to PKA-Cβ deletion. This supports the conclusion that PKA-Cα or PKA-Cβ have different downstream signaling networks.

**Table 7.**
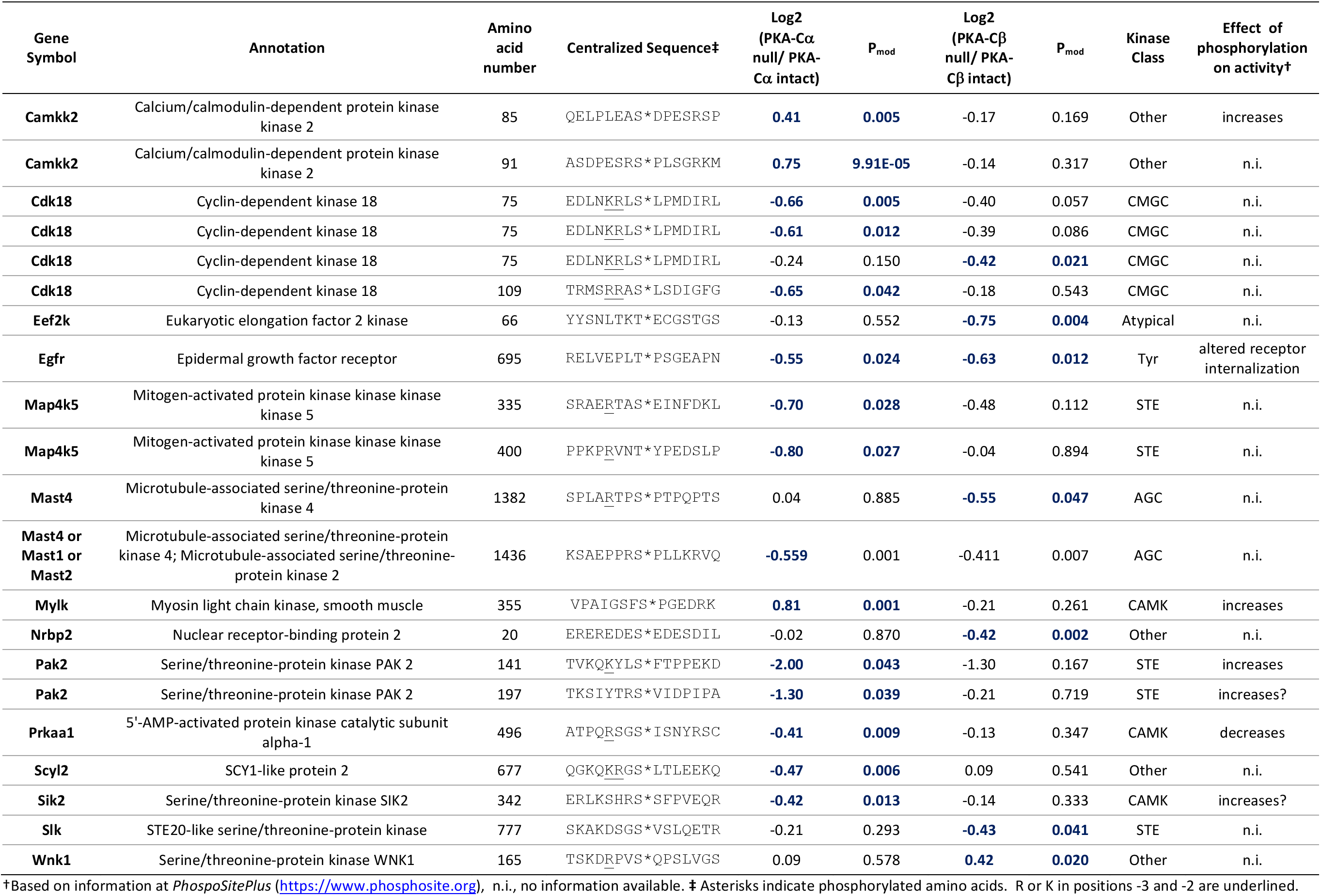
Protein kinases with altered phosphorylation in response to deletion of PKA-Cα or PKA-Cβ

### Differential regulation of cAMP signaling proteins by PKA-Cα and PKA-Cβ

One factor involved in localization of protein kinase A in the cell is its interaction with anchoring proteins such as AKAPs. **Table 8** shows altered phosphopeptides belonging to AKAP proteins or proteins with GO terms containing “cAMP” or “cyclic-AMP” to identify possible other AKAP interactors. Among the phosphoproteins in this list are Akap1, Akap12, cAMP phosphodiesterase 4C (Pde4c), cAMP phosphodiesterase 7A (Pde7a), AMP-activated protein kinase α-1 (Prkaa1), as well as two PKA regulatory subunits, RIα (Prkar1a) and RIIβ (Prkar2b). All of these (except for one site in Akap12) underwent changes in response to PKA-Cα deletion but not PKA-Cβ deletion. Thus, the phosphorylation evidence suggests that PKA-Cα interacts more extensively with components of AKAP complexes than does PKA-Cβ, at least with regard to phosphorylation.

**Table 8.** AKAPs and cAMP-associated proteins with altered phosphorylation in either PKA-Cα or PKA-Cβ-null cells

| UniProt ID | Gene Symbol | Annotation | Amino acid number | Centralized Sequence<sup>†</sup> | Log<sub>2</sub>(PKA-Cα null/ PKA-Cα intact) | P<sub>mod</sub> | Log<sub>2</sub>(PKA-Cβ null/ PKA-Cβ intact) | P<sub>mod</sub> |
| --- | --- | --- | --- | --- | --- | --- | --- | --- |
| O08715 | **Akap1** | A-kinase anchor protein 1, mitochondrial | 101 | TRQVRRRS*ESSGNLP | **-0.41** | 0.003 | 0.00 | 0.966 |
| Q9WTQ5 | **Akap12** | A-kinase anchor protein 12 | 22 | PAESDTPS*ELELSGH | **0.55** | 0.035 | -0.09 | 0.716 |
| Q9WTQ5 | **Akap12** | A-kinase anchor protein 12 | 583 | AEAEEGAT*SDGEKKR | **0.43** | 0.018 | 0.11 | 0.490 |
| Q9WTQ5 | **Akap12** | A-kinase anchor protein 12 | 584 | EAEEGATS*DGEKKRE | **0.64** | 0.008 | -0.02 | 0.908 |
| Q9WTQ5 | **Akap12** | A-kinase anchor protein 12 | 683 | KRARKASS*SDDEGGP | **0.51** | 0.011 | -0.11 | 0.514 |
| Q9WTQ5 | **Akap12** | A-kinase anchor protein 12 | 684 | RARKASSS*DDEGGPR | **0.52** | 0.019 | -0.20 | 0.305 |
| Q9WTQ5 | **Akap12** | A-kinase anchor protein 12 | 1292 | SNEEQSIS*PEKREMG | 0.40 | 0.061 | **0.49** | 0.026 |
| Q3UEI1 | **Pde4c** | cAMP-specific 3',5'-cyclic phosphodiesterase 4C | 301 | ELRRSSHT*SLPTAAI | **-0.43** | 0.034 | 0.26 | 0.173 |
| P70453 | **Pde7a** | High affinity cAMP-specific 3',5'-cyclic phosphodiesterase 7A | 58 | FETERRGS*HPYIDFR | **-0.44** | 0.020 | -0.40 | 0.031 |
| Q5EG47 | **Prkaa1** | 5'-AMP-activated protein kinase catalytic subunit alpha-1 | 496 | ATPQRSGS*ISNYRSC | **-0.41** | 0.009 | -0.13 | 0.347 |
| Q9DBC7 | **Prkar1a** | cAMP-dependent protein kinase type I-alpha regulatory subunit | 83 | DSREDEIS*PPPPNPV | **-0.90** | 0.001 | -0.11 | 0.599 |
| Q9DBC7 | **Prkar1a** | cAMP-dependent protein kinase type I-alpha regulatory subunit | 83 | DSREDEIS*PPPPNPV | **-1.05** | 1.34E-04 | -0.09 | 0.642 |
| Q9DBC7 | **Prkar1a** | cAMP-dependent protein kinase type I-alpha regulatory subunit | 83 | DSREDEIS*PPPPNPV | **-1.01** | 3.99E-04 | 0.05 | 0.816 |
| P31324 | **Prkar2b** | cAMP-dependent protein kinase type II-beta regulatory subunit | 112 | NRFTRRAS*VCAEAYN | **-1.38** | 0.001 | -0.29 | 0.402 |

<sup>†</sup>Asterisks indicate phosphorylated amino acids. R or K in positions -3 and -2 are underlined.

An important role of vasopressin-signaling in renal collecting duct cells is transcriptional regulation, particularly regulation of transcription of the *Aqp2* gene. **Table 9** shows transcription factors with altered phosphorylation in PKA-Cα-null or PKA-Cβ-null cells. Among all transcription factors, 241 phosphopeptides were quantified. There were 14 phosphopeptides that were substantially changed in abundance in 13 different transcription factor proteins. Most of the altered sites had proline in position +1 relative to the phosphorylated S or T, signifying altered phosphorylation by protein kinases in the cyclin-dependent kinase or MAP kinase families. Only one transcription factor underwent differential phosphorylation at a site with the (R/K)-(R/K)-X-p(S/T) motif, namely “cyclic AMP-dependent transcription factor ATF-7”, which showed a marked decrease in phosphorylation in the PKA-Cα-null but not the PKA-Cβ-null cells. Atf7 is a b-ZIP transcription factor that is inactive as a homodimer but can transactivate genes when heterodimerized with members of the AP-1 family including Jund (6), which underwent a decrease in phosphorylation. Beyond the transcription factors, various transcriptional co-regulators may participate in vasopressin-mediated regulation of transcription. One such protein is β-catenin, which shows increased phosphorylation at S552, a PKA target site (9), in response to vasopressin. Interestingly, S552 of β-catenin showed a marked decrease in phosphorylation in the PKA-Cα-null cells, but not the PKA-Cβ-null cells (**Table 3**). In general, as with other categories of proteins, changes in phosphorylation of transcription factors and their coregulators are different in PKA-Cα-null or PKA-Cβ-null cells, again supporting the conclusion that the two PKA catalytic subunits fulfill different cellular functions.

**Table 9.** Phosphopeptides from transcription factor proteins that underwent changes in abundance with PKA-Cα and/or PKA-Cβ deletion

| UniProt ID | Gene Symbol | Annotation | Amino acid number | Centralized sequence<sup>†</sup> | Log2 (PKA-Cα null/ PKA-Cα intact) | P<sub>mod</sub> | Log2 (PKA-Cβ null/ PKA-Cβ intact) | P<sub>mod</sub> | TF family |
| --- | --- | --- | --- | --- | --- | --- | --- | --- | --- |
| Q3US59 | **Atf7** | Cyclic AMP-dependent transcription factor ATF-7 | 326 | TGGRRRRT*VDEDPDE | **-1.19** | 0.039 | -0.60 | 0.266 | bZIP |
| Q61103 | **Dpf2** | Zinc finger protein ubi-d4 | 142 | DPRVDDDS*LGEFPVS | -0.44 | 0.061 | **-0.59** | 0.017 | zf-C2H2 |
| Q8K5C0 | **Grhl2** | Grainyhead-like protein 2 homolog | 2 | _____MS*QESDNNK | **0.42** | 0.038 | -0.11 | 0.539 | CP2 |
| P15066 | **Jund (or Jun)** | Transcription factor jun-D (or jun) | 100 | LGLLKLAS*PELERLI | **-0.69** | 0.002 | 0.07 | 0.720 | bZIP |
| Q2VPU4 | **Mlxip** | MLX-interacting protein | 9 | AADVFMCS*PRRPRSR | -0.03 | 0.897 | **-0.68** | 0.010 | bHLH |
| P40645 | **Sox6** | Transcription factor SOX-6 | 439 | KTAEPVKS*PTSPTQN | **1.40** | 0.002 | 0.13 | 0.708 | HMG |
| P40645 | **Sox6** | Transcription factor SOX-6 | 454 | LFPASKTS*PVNLPNK | **1.19** | 0.007 | -0.10 | 0.783 | HMG |
| Q9WTN3 | **Srebf1** | Sterol regulatory element-binding protein 1 | 96 | KVTPAPLS*PPPSAPA | **-0.80** | 0.024 | 0.22 | 0.500 | bHLH |
| Q9JM73 | **Srf** | Serum response factor | 220 | TCLNSPDS*PPRSDPT | **0.41** | 0.007 | -0.05 | 0.710 | SRF |
| P63058 | **Thra** | Thyroid hormone receptor alpha | 12 | PSKVECGS*DPEENSA | **0.61** | 0.006 | -0.24 | 0.208 | TH |
| P62960 | **Ybx1** | Nuclease-sensitive element-binding protein 1 | 174 | EKNEGSES*APEGQAQ | **-0.47** | 0.020 | -0.05 | 0.792 | CSD |
| Q3UQ17 | **Zbtb16** | MCG3834 | 307 | EESGEQLS*PPVEAGQ | -0.44 | 0.236 | **-1.04** | 0.012 | ZBTB |
| Q8BIQ6 | **Zfp947** | MCG23335 | 283 | FTQKSSLS*IHQMYHT | **0.50** | 0.044 | 0.07 | 0.768 | zf-C2H2 |
| O35615 | **Zfpm1** | Zinc finger protein ZFPM1 | 681 | RGSEGSQS*PGSSVDD | **0.68** | 0.030 | -0.06 | 0.843 | zf-C2H2 |

<sup>†</sup>Asterisks indicate phosphorylated amino acids. R or K in positions -3 and -2 are underlined.

## DISCUSSION

Based on the observations in this paper, we conclude that PKA-Cα and PKA-Cβ do not have redundant functions. Indeed, while some overlap exists between the two in terms of phosphorylation targets, large differences were seen at a whole phosphoproteome level. PKA-Cα-null cells showed decreased phosphorylation dominated by sites with the classical PKA motif, viz. (R/K)-(R/K)-X-p(S/T), while only very few of the decreased phosphorylation sites in the PKA-Cβ-null cells contained this motif. However, in vitro incubation with recombinant PKA-Cα and PKA-Cβ resulted in phosphorylation of virtually identical sites, with a predominance of the (R/K)-(R/K)-X-p(S/T) motif. A key question is, “If the catalytic regions of PKA-Cα and PKA-Cβ have nearly identical in vitro target specificities, why is there such a difference in phosphorylation targets in the intact cells?”. The question was addressed by the bioinformatics analysis of phosphorylation targets, showing that completely different *Gene Ontology Cellular Component* terms are associated with the two sets of phosphorylation targets. Specifically, PKA-Cα targets were largely related to cell membranes and membrane vesicles, while PKA-Cβ targets were related to the actin cytoskeleton and cell junctions, indicating that PKA-Cα and PKA-Cβ interact with different sets of proteins in cells. This difference could arise from differences in the non-catalytic regions of the two kinases that result in a different set of local interactions with potential substrates.

One possibility is that the two catalytic subunits could interact with different AKAPs and/or PKA regulatory subunits (15). Phosphoproteomic results point to a preferential association of PKA-Cα with the regulatory R2 β subunit (Prkar2b), which showed a marked decrease at S112 in the PKA-Cα null cells but not the PKA-Cβ null cells **(Table 8)**. The second possibility is that there could be differential interactions with so-called C-kinase anchoring proteins (C-KAPs) (25). The third possibility is that there may be a difference in protein-protein interactions with PDZ domain-containing proteins (1). PKA-Cα contains a class-1 carboxy terminal PDZ-ligand motif (–TxF), while PKA-Cβ, ends with –CxF which does not conform to a PDZ ligand motif. In previous studies (3), we showed that vasopressin treatment of native inner medullary collecting duct cells results in increased phosphorylation of several PDZ domain-containing proteins (connector enhancer of kinase suppressor of Ras 3 (Cnksr3), PDZ domain-containing protein GIPC1 (Gipc1), multiple-PDZ domain protein (Mpdz), and PDZ and LIM domain protein-5 (Pdlim5). That paper showed that proteins with class I COOH-terminal 3-amino acid motifs are more likely to show increases in phosphorylation in response to vasopressin than proteins without the motif.

In addition to phosphorylation differences, PKA-Cα and PKA-Cβ deletions resulted in many differences in what proteins underwent changes in total protein abundances. Thus, in terms of regulation of protein abundances, PKA-Cα and PKA-Cβ proteins appear to have non-redundant functions. For example, AQP2 was markedly decreased in PKA-Cα-null cells but not in PKA-Cβ-null cells. This result suggests that PKA-Cα is the predominant catalytic subunit responsible for the regulation of AQP2 abundance. Similarly, complement component C3 was markedly decreased in PKA-Cα-null cells but not in PKA-Cβ-null cells. In contrast, the abundance of mucin-4 (Muc4) was substantially decreased in PKA-Cβ-null cells but not in PKA-Cα-null cells. Prior studies in mpkCCD cells have shown that, in the PKA double knockout cells AQP2, C3, and Muc4 mRNA and protein levels were markedly decreased (9).

### Data resource

In addition to the scientific findings highlighted above, this paper provides added value in the form of two web resources that allow users to interrogate data from this paper. These resources have been included with other phosphoproteomic data on the *Kidney Systems Biology* Project website (https://hpcwebapps.cit.nih.gov/ESBL/Database/).

## ACKNOWLEDGEMENTS

This work was funded by the Division of Intramural Research, National Heart, Lung, and Blood Institute (projects ZIA-HL001285 and ZIA-HL006129, MAK). Karim Salhadar was a member of the Biomedical Engineering Student Internship Program (BESIP, Robert Lutz, Director) supported by the National Institute for Biomedical Imaging and Bioengineering (June-August 2018). Mass spectrometry utilized the NHLBI Proteomics Core Facility (M. Gucek, Director). Supplemental data are deposited at https://hpcwebapps.cit.nih.gov/ESBL/Database/Supplemental_Data_PKA_sKO/.

## Author Contributions

V.R., and M.A.K., developed the concept and designed the experiments. V.R., K.S, E.P., and C-R.Y., performed the experiments. V.R., K.L., and M.A.K., analyzed the data. V.R., K.L., C-R.Y., and M.A.K interpreted the results of the experiments. V.R., K.S., K.L., and M.A.K., drafted the manuscript and prepared figures. All authors edited, revised, and reviewed the manuscript.

## Competing interests

The author(s) declare no competing interests.

## Data availability

The mass spectrometry proteomics data have been deposited to the ProteomeXchange Consortium via the PRIDE partner repository with the dataset accession number PXD015050 (URL for reviewer is https://www.ebi.ac.uk/pride/archive/projects/PXD015050.

## Notes

https://hpcwebapps.cit.nih.gov/ESBL/Database/Supplemental_Data_PKA_sKO/

### Competing Interest Statement

The authors have declared no competing interest.

